# Stochastic Modeling of BMP Heterodimer-Receptor Interactions Shows Emergence of Low-Pass Filtering Behavior

**DOI:** 10.1101/2025.10.17.677951

**Authors:** Nissa J. Larson, Aasakiran Madamanchi, Linlin Li, David M. Umulis

**Affiliations:** Weldon School of Biomedical Engineering, Purdue University, West Lafayette, IN; School of Information, University of Michigan, Ann Arbor, MI

## Abstract

In developing tissues, signal transduction from morphogen gradients conveys positional information to cells, resulting in cell specification and differentiation. One such morphogen is bone morphogenetic protein (BMP), of the TGF-β superfamily, whose signaling network is highly conserved across many species. In *Danio rerio* (zebrafish), this signaling pathway directs dorsoventral axis formation during early embryogenesis. Many of the molecules that play a role in this network are well-understood; however, the mechanisms through which they achieve noise attenuation and gradient robustness have not been fully defined. Specifically, the heterodimer-heterotetramer complex has been shown to be required for signal transduction[1], but current understanding and modeling of the BMP membrane receptors at this stage has not given any insight into evolutionary drivers of the requirement. In this study, we develop a stochastic model of receptor oligomerization with the published reports of binding kinetics of BMP ligand-receptor interactions to mechanistically assess zebrafish phenotype variability related to the distributions of noise and stochasticity. We can also analyze time-dependent signaling and frequency metrics that are not available in traditional, deterministic modeling. Fast Fourier Transform and cumulative energy spectral density visualization show that the heterodimer-heterotetramer complex may function as part of a low-pass filter mechanism in the dorsal-ventral axis formation process, specifically tuned to the noise of the system. Under dynamic conditions such as the mid-blastula transition (MBT), wherein the morphogen gradient rapidly changes shape, established metrics of noise and information transduction, such as coefficient of variation and mutual information, overlook important temporal effects that may be particularly relevant during development. As the BMP signaling pathway is highly conserved and has been implicated in human bone growth and wound healing, its study in simpler systems stands to accelerate our comprehension of BMP network structure and molecular mechanisms with potential application in regenerative medical studies.

## Introduction

### BMP signaling in embryogenesis requires heterodimer-heterotetramer complex

Embryonic patterning, the organization of initially homogeneous cells into differentiated tissues, is an integral process in normal embryonic development. In living systems, this process is mediated through the cellular interpretation of morphogen signals. Morphogens, typically chemical ligands, are substances that form a spatial gradient along which they induce a distinct set of concentration-dependent cellular responses[2], [3], [4], [5]. In many vertebrate systems, the key signaling pathways involved in embryonic patterning have already been characterized; in zebrafish (*Danio rerio*) dorsoventral patterning is regulated by the temporally precise induction of bone morphogenetic protein (BMP) signaling[6], [7]. Despite the extensive study of BMP signaling transduction, a systematic understanding of the design principles underlying the evolution of such signaling mechanisms remains lacking. In this study, we aim to determine the attributes of the complex signaling pathway that contribute to robustness in the face of noise and perturbation, allowing the system to uphold the standard for axis formation at this stage and throughout the life of the organism.

The BMP signaling system is a primordial, evolutionarily conserved pathway which directs patterning and dorsoventral axis formation in both vertebrate and invertebrate systems. BMP ligand signals through an oligomeric complex consisting of a dimeric ligand and tetrameric set of receptors. The tetrameric receptor complex requires two Type I receptors (BMPR1 and/or Acvr1) and two Type II receptors. Within a ligand-bound tetrameric receptor complex, the Type II receptors, which are constitutively active, phosphorylate and activate the Type I receptors[8]. The kinase function of the activated Type I receptor initiates intracellular signaling by phosphorylating cytoplasmic R-Smads (Smads 1/5/9). Phosphorylated R-Smads bind with a co-Smad (Smad4) to form an active Smad complex, which accumulates in the nucleus due to ongoing nucleocytoplasmic shuttling[9]. In the nucleus, active Smad complexes interact with transcriptional machinery to regulate expression of BMP target genes[8], [10].

Experimental evidence from both *Drosophila* and zebrafish indicates that during dorsoventral axis formation, the physiologically relevant ligand-receptor oligomeric complex consists of the BMP2/7 ligand heterodimer and a heterotetrametric receptor complex with one BMPR1 receptor, one Acvr1 receptor, and two Type II receptors[7]. Experimental evidence suggests that other BMP homodimer and homotetramer complexes are abundant during this developmental window, so it remains unclear why the BMP heterodimer-heterotetramer remains the obligate ligand-receptor signaling complex during dorsoventral axis formation across millions of years of evolution.

In multiple developmental systems, BMP-mediated embryonic patterning appears to be tightly temporally and spatially coordinated[6], [11], [12], [13], [14]. In zebrafish, zygotic BMP expression first occurs after the mid-blastula transition (MBT) at 3 hours post fertilization[15], [16], [17], [18]. Initially, BMP expression is ungraded, but interactions with Chordin and other regulators lead to the rapid formation of a gradient[11], [13]. Imaging and modeling approaches suggest that BMP-induced p-Smad signaling remains ubiquitous through the Sphere stage (4 hpf); at 30% epiboly, the gradient appears, with dorsal p-Smad being negligible and ventral p-Smad signaling being elevated[14]. Ventral p-Smad signaling continues to increase, sharpening the gradient, through the shield stage (6 hpf). The speed of the morphogen gradient formation is necessary as the correct BMP gradient is required at the onset of gastrulation to pattern rostral DV cell fates[6]. BMP autoregulation begins at 5.7 hpf and helps to shape a dynamically evolving BMP gradient which is required for the specification of more caudal DV cell fates at progressively later intervals during gastrulation[6], [11]. A temporally evolving BMP signaling gradient appears to regulate neuronal subtype identity in the dorsal neural tube as well[12]. Previously, our research concentrated on examining the extracellular regulation of BMP patterning formation during the blastula and gastrula stages of zebrafish embryonic development.[11], [19], [20]. We found that the source-sink mechanism and advection caused by the cell flow during epiboly plays a role in maintaining the BMP gradient during patterning. It remains unclear what role ligand-receptor oligomerization plays in gradient formation and interpretation. Prior attempts to use computational approaches to understand the role of BMP receptor oligomerization have been limited to steady-state analyses. Antebi et al. modeled a BMP system with a simplified receptor oligomerization schema using deterministic approaches to understand how the simultaneous presence of multiple ligands is interpreted in cellular systems[21] [22]. Previous work from our group has employed stochastic modeling of a single ligand dimer BMP system to analyze the effect of an extracellular regulator, Crossveinless-2 (Cv-2), on noise propagation[23], and kinetic modeling of BMP tetramer formation to analyze the theorized favorability of the heterodimer-heterotetramer complex[24]. Outside of the BMP system, researchers have begun to draw on the field of information theory to quantitatively calculate the amount of ‘positional information’ transmitted by a given morphogen[25], [26]. This methodological advance brings forth the opportunity to understand the fundamental information transduction properties of different morphogen interpretation systems. In this paper, we use a stochastic computational modeling approach to understand the information transduction properties of bone morphogenetic protein (BMP) receptor oligomerization in developmental contexts. Specifically, we developed a kinetic-based stochastic model of the complete BMP ligand-receptor oligomerization system and applied this model to simplified 1-D embryos with experimentally informed morphogen gradients.

### Stochastic model structure considers 51 possible species and outcomes

Formation of BMP ligand-receptor signaling complexes occurs through a dynamic series of physiological, reversible binding events between receptors and ligand-receptor oligomers. A diagram of all possible reaction steps in the oligomerization process for a single ligand dimer is shown in Figure 1. All three possible ligand dimers, BMP2/2, BMP2/7, and BMP7/7 are present and ‘compete’ for available receptors at the same time. The complete model tracks 51 possible ligand-receptor oligomeric complexes and includes nine complete ligand-receptor tetramer complexes, as sub functionalization and homodimer contribution have been suggested to be required[24]. The forward and reverse reaction rates for each binding event are drawn from published experimental data in the structural biology literature [supplemental]. This model development process was previously used to model *Dpp* oligomerization in *Drosophila*[23].

**Figure 1.**
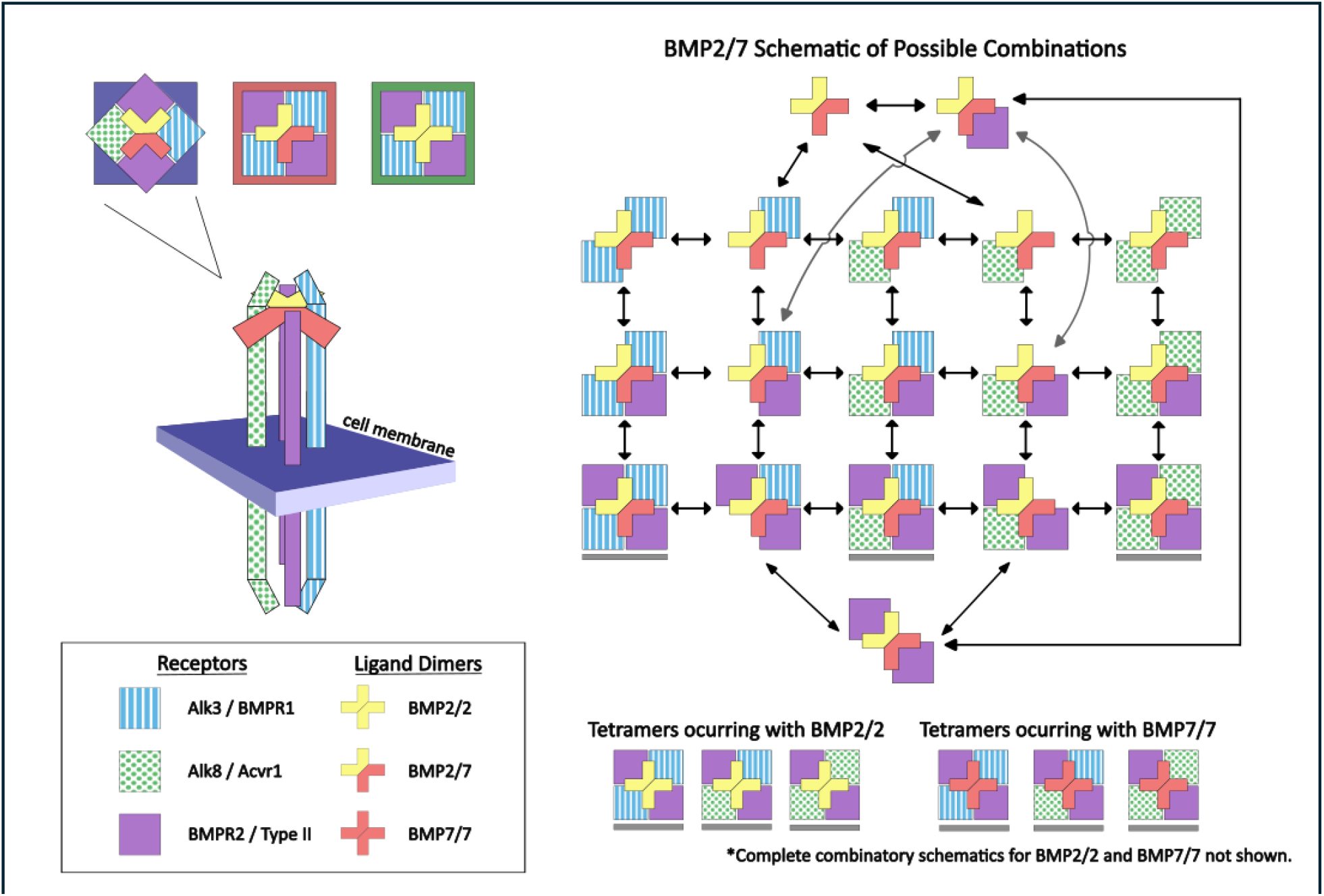
BMP ligand-dimer receptor-tetramer oligomerization reactions. BMP signaling occurs through an oligomeric signaling complex, which necessarily consists of a ligand dimer (red/gold cross) bound to a receptor tetramer. The receptor tetramer requires two Type I receptors (BMPR1 in blue and/or Acvr1 in green) and two Type II receptors (purple square). Oligomeric signaling complexes are formed through reversible reactions (double-sided arrows). The reaction schematic is shown for BMP2/7. All combinations of receptors can also occur for BMP2/2 and BMP 7/7; however, some combinations are much less likely than others (see the supplementary material for reaction kinetics). The three most abundant complexes are highlighted on the top left. They are BMP2/7-Acvr1-BMPR1-TypeII_2_ in blue, BMP2/7-BMPR1_2_-TypeII_2_ in red, and BMP2/2-BMPR1_2_-TypeII_2_ in green. In all following figures, data correlates with these highlighted colors.

All data come from simulations conducted with a 1:1:2 stochiometric ratio of BMPR1, Acvr1, and the Type II receptor. The initial values are 3500 molecules of each Type I bmp receptor and 7000 molecules of the Type II bmp receptor unless otherwise indicated[27]. Simulation volume was 1.25 nanoliters, an estimate for immediate volume surrounding cell membrane[28]. BMP ligand dimer influx and initial receptor binding is modeled as a zeroth-order reaction, and quantities are simulated as constant availability as previously published[24].

The stochastic model was constructed within the StochKit2 stochastic simulation framework, which implements tau-leaping and Gillespie’s stochastic simulation algorithm in C++ to facilitate rapid simulation of stochastic trajectories in large biochemical networks. The StochKit2 framework is accessed through GillesPy, an open-source Python wrapper which facilitates cluster-implementation and enables the screening of a larger parameter space[29].

Positional information is calculated by taking the mutual information of cellular position and tetramer or pSmad levels. This approach provides a representation of information available to cells through local signaling. Positional information was calculated using the ‘direct method’ with an input of the gradient of the available signaling molecule[26]. Fourier analysis provides a readout of signaling frequencies contained within a signal. Two signals in the frequency domain can be more effectively compared in their informational availability on a time-frequency basis.

In addition to simulating steady-state morphogen fields, we also simulated the dynamic ligand gradients in a virtual blastula formation experiment. Specifically, we simulate 1-D morphogen fields for the first three hours after MBT. One hundred replicates are used to generate dynamic positional information data.

## Results

### Parameter screening validates model structure

Although we model all 51 possible ligand-receptor oligomers, including nine complete tetrameric receptor oligomers (Figure 1), in this paper, we only display data for the three most abundant receptor tetramers. In all figures, data for BMP2/2-(BMPR1)_2_-(Type II)_2_ is displayed in green, data for BMP2/7-(BMPR1)_2_-(Type II)_2_ is displayed in red, and data for BMP2/7-Acvr1-BMPR1-(Type II)_2_ is displayed in blue.

To validate our stochastic model, we demonstrate that a deterministic ODE model of the BMP ligand-receptor oligomerization reactions produces data consistent with the average result, across multiple runs, of our stochastic model. In Figure 2A, we show the steady-state concentration of each candidate receptor tetramer in deterministic simulations with ligand concentrations ranging from 0.003 nM to 3 nM. Physiologically relevant ligand levels are considered to be 0.01 to 1 nM based on previously published estimates[23]. In Figure 2B, we show corresponding results from stochastic simulations. Specifically, stochastic simulations were run at each ligand concentration for 48 hours to observe and quantify the long-term dynamics of the ligand-receptor oligomerization. The mean ± SD for each candidate receptor tetramer over the second 24 hours (to ensure steady-state values) of each simulation are displayed in Figure 2B. Stochastic and deterministic simulations show similar steady-state receptor tetramer levels at each ligand concentration.

**Figure 2.**
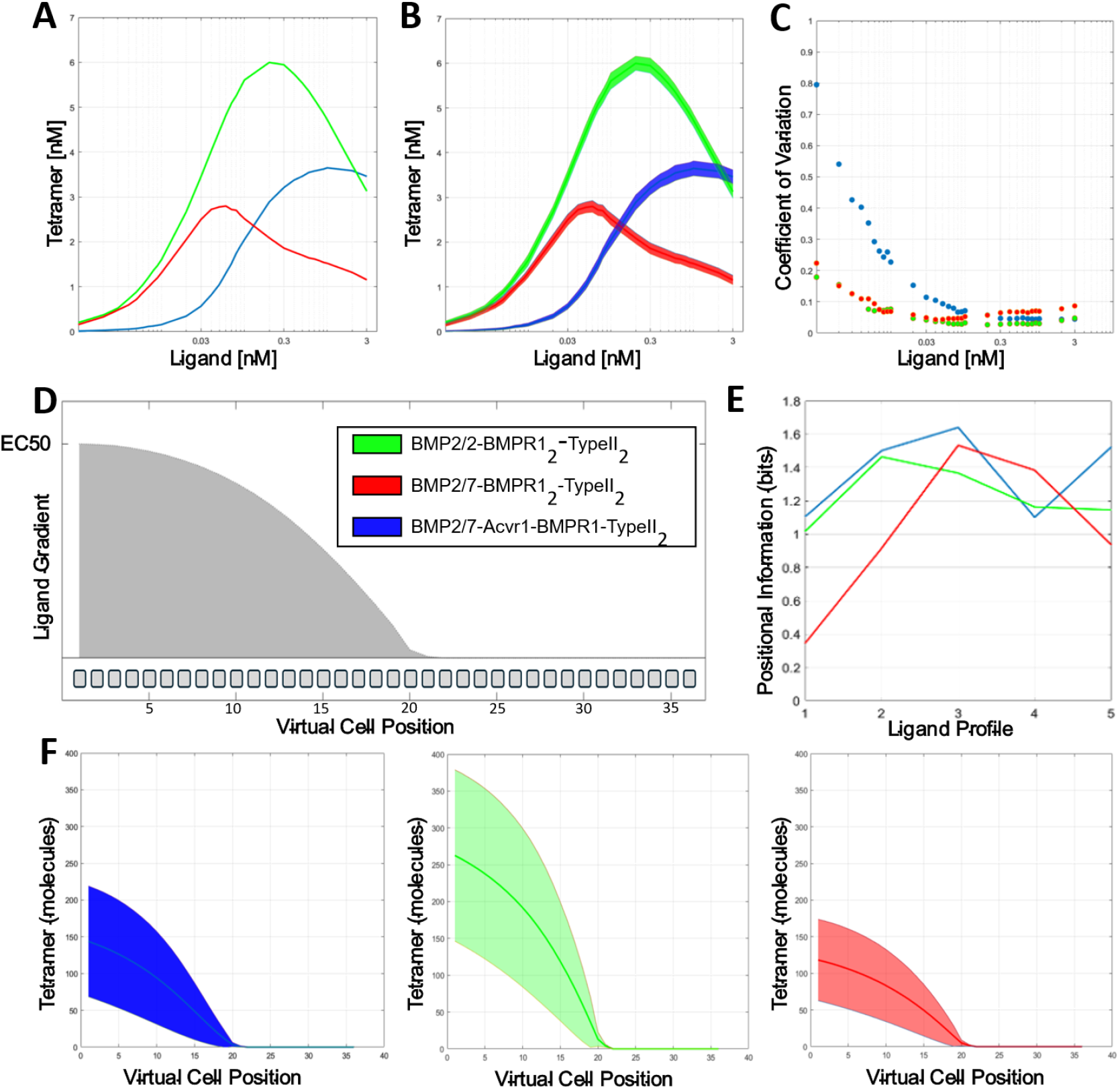
Noise and positional information of BMP receptor-tetramer signaling complexes. **A**. Deterministic simulation of steady-state levels for each candidate receptor tetramer. **B**. Stochastic simulated mean SD of steady-state tetramer levels over 24 hours. **C**. Coefficient of variation (SD/mean) of each receptor tetramer at various ligand levels. **D**. Schematic for 1-D morphogen field simulation showing EC50 normalized ligand gradient over a field of 36 cells. **E**. Positional information of each receptor tetramer over a gradient beginning at a given ligand level for five gradients. **F**. Densiometric visualization of 240 hours of steady state levels of each tetramer in the morphogen field simulation. Positional information (PI) is labeled. In all figures, data for BMP2/2-BMPR1_2_-TypeII_2_ shown in green, BMP2/7-BMPR1_2_-TypeII_2_ shown in red, and BMP2/7-Acvr1-BMPR1-TypeII_2_ shown in blue.

We follow with an analysis of the noise generated in the oligomerization process by each receptor tetramer at physiologically relevant ligand levels. In Figure 2C, the coefficient of variation (CV), a mean-normalized measure of noise, is calculated from the simulation data in 2B. At lower ligand levels, the heterodimer-heterotetramer shows a higher coefficient of variation compared to the other candidate receptor tetramers. However, as the ligand level increases, the coefficient of variation of heterodimer-heterotetramer decreases significantly. This result suggests that the heterodimer-heterotetramer has strong capabilities in noise reduction.

Although noise can be modeled at individual ligand levels, positional information can only be calculated across a ligand gradient. We generated a hypothetical 1-D virtual zebrafish embryo (Figure 2D) that represents thirty-six cells along the leading margin of epiboly, where BMP patterns cell fate, while a data-derived morphogen gradient is applied between the ligand source (in cell 1) and the end of the embryo in cell 36. The absolute values of BMP ligand gradients in gastrulating zebrafish embryos have not yet been experimentally determined, so we use ligand gradients scaled from 3E-2, 3E-1.6, 3E-1.2, 3E-0.8, 3E-0.4, and 3E-0 nM to test positional information at different ligand levels. In Figure 2E, we show the positional information obtained from these stochastic 1-D virtual embryo simulations. The heterodimer-heterotetramer shows greater positional information transmission in the virtual embryos with lower ligand levels, but lower positional information than other receptor tetramers in virtual embryos with higher physiological ligand levels.

Results in Figure 2B-E indicate that the relative noise and information properties of each receptor tetramer vary with different ligand levels. In part, this is because different ligand levels reflect different receptor level saturation for each receptor tetramer candidate. At a lower ligand level, the heterodimer-heterotetramer is in the optimal part of its dynamic ligand range, while at higher ligand levels, the other candidate tetramers have the advantage. To isolate the specific noise and information properties of each receptor tetramer, we customize the morphogen gradient in a 1-D virtual embryo setup (schematic in Figure 2D) by scaling the ligand gradient from the EC50 level of each receptor. Half Maximal Effective Concentration (EC50) is a commonly considered, biologically-relevant concentration, at which a half peak response is recorded. In essence, each tetrameric complex has the greatest opportunity to provide positional information in a simulation run with its own EC50 ligand concentration. The three panels in Figure 2F show densitometric visualization of 240 hours of simulated steady-state receptor tetramer levels in each cell of the customized virtual embryo, each with its respective EC50 ligand concentration, obtained through data analysis of Figure 2B. The positional information, calculated using previously published methods, is also listed[25], [26]. Contrary to our expectation, the heterodimer-heterotetramer provides less positional information in this neutral test than the other candidate receptor tetramers.

### Frequency analysis of modeled receptor response shows heterodimer-heterotetramer advantage

To further understand the noise and information propagation properties of each receptor tetramer, we sought to isolate the relative contributions of high and low frequency noise. Using the Fast Fourier Transformation (FFT), a frequency analysis of fluctuations in receptor tetramer levels was conducted at steady state in simulations with EC50 ligand levels. Normalized cumulative progressions of energy spectral density (ESD) for each candidate tetramer receptor are shown in Figure 3A, while the component of the normalized ESD from various frequency bins is shown in Figure 3B. Together, these graphs indicate that the heterodimer-heterotetramer exhibits relatively fewer low-frequency oscillations (> 10^-4^ Hz) and relatively greater high-frequency oscillations compared to the other candidates. To confirm the Fourier analysis, we visually inspected simulated per-second trajectories of each candidate tetramer receptor. In Figure 3C, we display six hours of steady state time series data in mean-normalized, zero-centered form for each candidate receptor tetramer. Tetramer levels that are more than a standard deviation above or below the mean steady state level for that receptor tetramer, are considered departures or errors. In Figure 3C, the errors for each receptor tetramer are highlighted. Compared to the other candidate tetramer receptors, the heterodimer-heterotetramer has a similar total time outside of one standard deviation from the mean; however, it has fewer long-lasting errors. Note that in the visualized time series data (Figure 3D), both BMP2/2-(BMPR1)_2_-(Type II)_2_ (green) and BMP2/7-(BMPR1)_2_-(Type II)_2_ (red) have errors spanning greater than half an hour. In contrast, the heterodimer-heterotetramer has no errors greater than thirty minutes in duration. To demonstrate that this comparison is representative of the overall data, we tabulated the incidence of errors of various lengths across 240 hours of steady-state simulation. Errors are only tabulated from the ventral (higher-BMP-expressing) half of the simulated morphogen fields to filter away the increased noise in the low abundance half of the simulated morphogen field (Figure 2E). Longer lasting departures or errors are potentially maladaptive in biological systems under autoregulation, as in gastrulation, or under tight time constraints, as in blastula formation. Therefore, reliance on the heterodimer-heterotetramer complex for signal transduction during these few hours of embryogenesis may drive higher accuracy in downstream cell differentiation through fewer low-frequency fluctuations.

**Figure 3.**
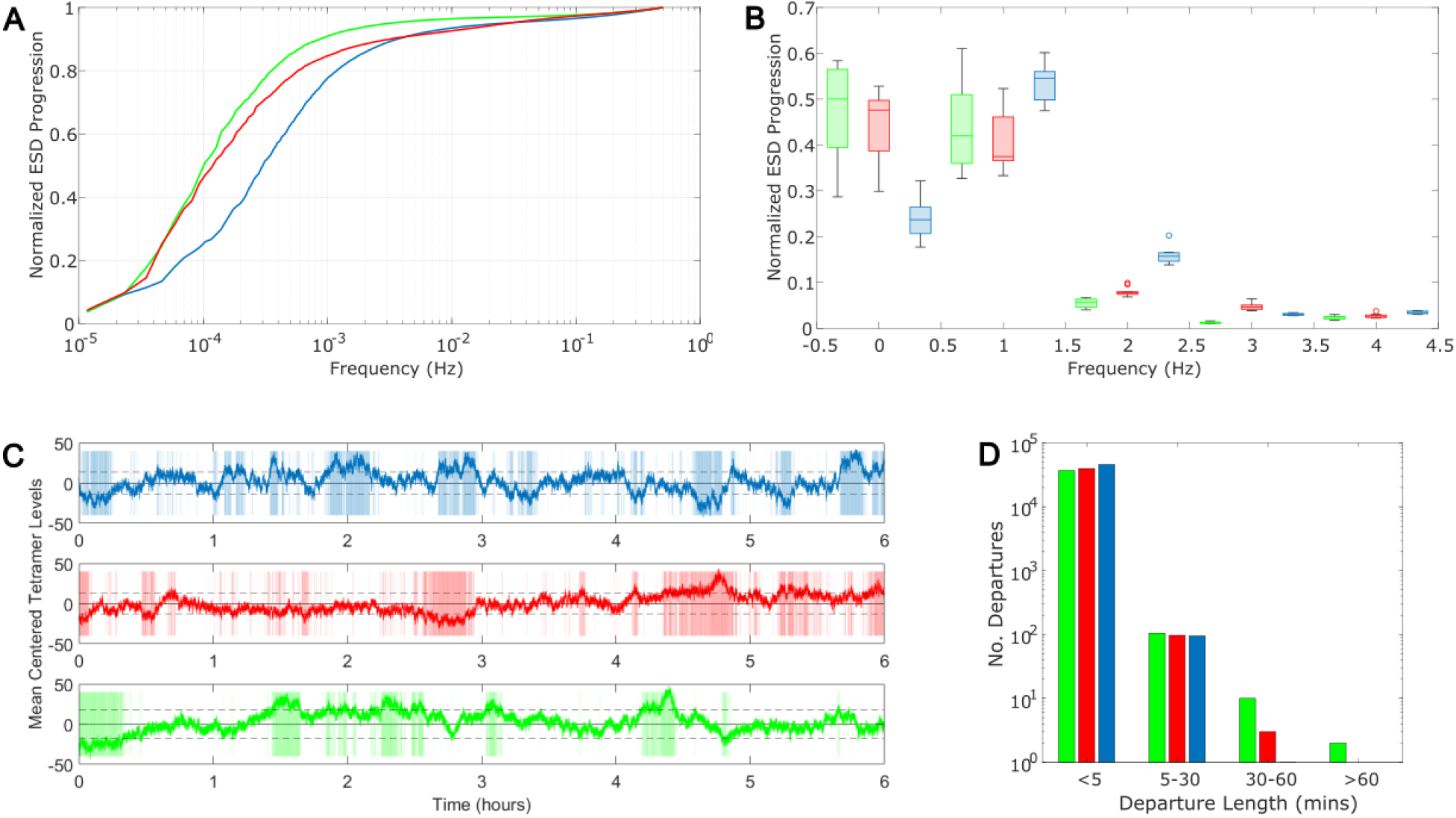
High vs low frequency noise of BMP receptor tetramers. **A**. Normalized cumulative energy spectral density (ESD) levels for each candidate tetramer receptor as calculated by Fast Fourier Transformation (FFT). **B**. Relative contribution of each frequency band to the ESD **C**. Mean-normalized zero-centered levels of each receptor tetramer over six hours of steady state simulation. Illustrative data: areas where tetramer levels are outside of one standard deviation (errors) are highlighted. **D**. The number of departures of a given length per 240 hours of steady state simulation. In all figures, data for BMP2/2-BMPR1_2_-TypeII_2_ shown in green, BMP2/7-BMPR1_2_-TypeII_2_ shown in red, and BMP2/7-Acvr1-BMPR1-TypeII_2_ shown in blue.

### Sensitivity analysis conveys receptor ratios have major impacts on system behavior

The data in Figures 2F and 3 were generated from stochastic simulations of our model using a total quantity of 14,000 BMP receptors. This receptor quantity was based on estimates of 10,000-20,000 TGF-β receptors per cell[27]. There is limited evidence on the relative representation of the BMP receptor subtypes during gastrulation. In these experiments, we have assumed a stochiometric 1:1:2 ratio of BmpR1, AcvR1 and Type II receptors based on requirement for two Type I and two Type II receptors in every receptor tetramer. To assess the robustness of our findings against different BMP receptor availabilities, we conducted a series of single-factor sensitivity analyses for each BMP receptor subtype. Single-factor sensitivity was required as the computational expense of a global sensitivity analysis with stochastic simulations was prohibitive. Specifically, we individually modulated the number of Acvr1, BMPR1, and the Type II receptors in the 1D virtual embryo system in Figure 2F. We examined the effects of both decreased receptor availability (0.25-, 0.5-, and 0.75-fold expression of the modulated receptor) and increased receptor availability (1.5-, 2-, 4-, 8-, 10-, and 15-fold expression of the receptor in question). As mentioned in the previous section, the ligand gradient was scaled to the EC50 of each receptor tetramer in each model condition to minimize the effect of differences in the dynamic ligand range of each receptor tetramer. Model conditions in which the dynamic range is too small to differentiate the EC50 from the lowest point in the ligand gradient are not shown.

In Figure 4A, we present the positional information for each receptor tetramer in simulations with varying BMP receptor levels. In the second panel, we observe that increased BMPR1 availability allows for greater information transmission by both homomeric receptor tetramers (red and green). The heterodimer-heterotetramer also gains advantages in positional information transmission with upregulation of Acvr1; with 8-, 10-, and 15-fold upregulation of Acvr1, the heterodimer-heterotetramer has more positional information than the other receptor tetramers. Interestingly, BMP2-(BMPR1)_2_-(Type II), (green) is largely unaffected by increased expression of Acvr1, whereas information transmission via BMP2/7-(BMPR1)_2_-(Type II)_2_ (red) decreases as Acvr1 increases. This divergent response is likely to arise from the difference in relative affinity for Type I receptor subtypes between the BMP2 and BMP7 ligand monomers.

**Figure 4.**
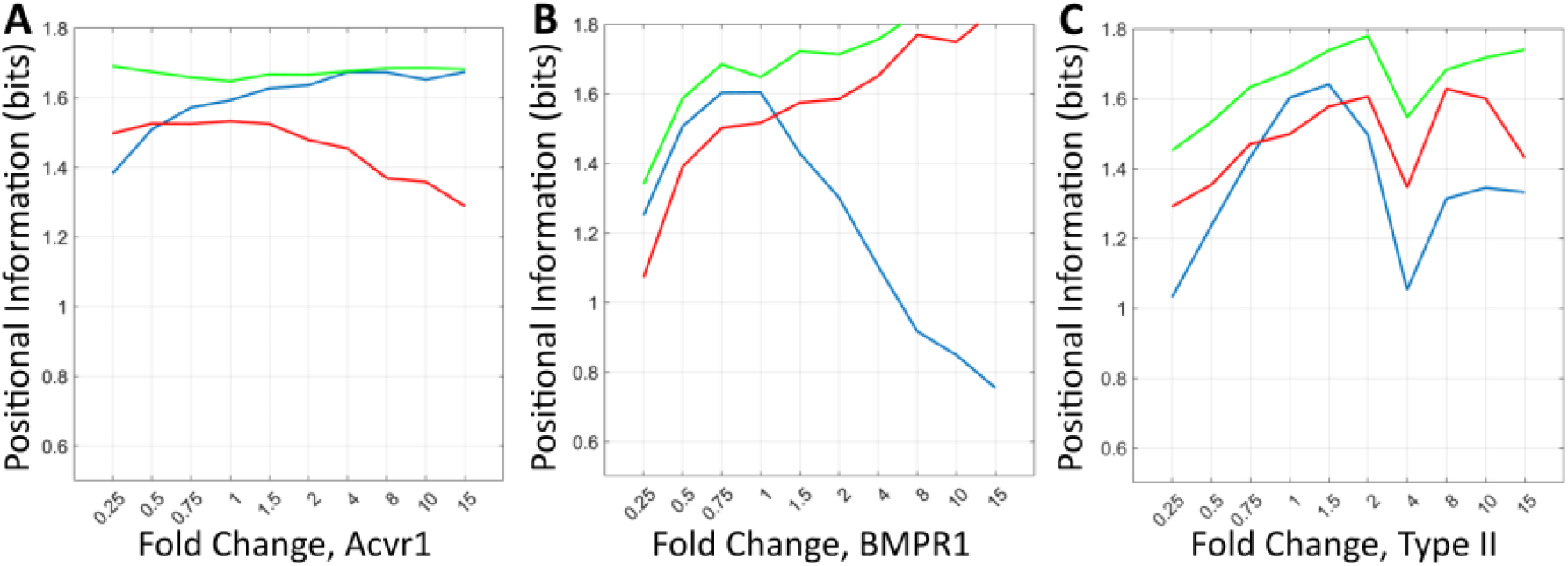
Effect of BMP receptor levels on positional information and error duration of each receptor tetramer. Levels of **A**. Acvr1, **B**. BMPR1, and **C**. Type II receptors are individually modulated, and one-dimensional morphogen fields are simulated for each receptor tetramer. In all figures, data for BMP2/2-BMPR1_2_-TypeII_2_ shown in green, BMP2/7-BMPR1_2_-TypeII_2_ shown in red, and BMP2/7-Acvr1-BMPR1-TypeII_2_ shown in blue.

Computational modeling efforts often rely on key assumptions in areas with limited experimental data, and this work stands to be validated or enhanced with the collection of more data from which to draw. This acknowledgment highlights both the strength and weakness of computational modeling, in that infinite theoretical data can be generated, which can drive biological phenomena questions, but the uncertainty of motivating factors can bring about skepticism. This bulk of this work utilizes an equal ratio of type I to type II receptors, with equal ratios of subtypes within them, as he simplest reasonable explanation for a tetrameric complex receptor forming network.

### Heterodimer-heterotetramer displays improved functionality when paired with intracellular phosphorylated Smad acting as a low-pass filter

Oligomeric BMP signaling complexes induce cellular responses through the phosphorylation and nuclear localization of R-Smads[8]. Downstream signal transduction is known to have essential roles in noise filtering and information transmission in cellular signaling systems[30], [31]. To understand how signal processing through downstream Smad signaling might affect cellular interpretation of BMP oligomerization, we integrated a simple model of Smad activation.

Zebrafish gastrulation requires the rapid interpretation of a dynamically evolving BMP ligand gradient. In the early blastula, ubiquitous ungraded BMP ligand expression rapidly evolves into a gradient after midblastula transition (MBT). To understand how the receptor tetramers respond to a dynamic ligand gradient in this developmental context, we constructed a simplified virtual blastula formation simulation in which we simulate 100 1-D morphogen fields for the first three hours after MBT. There is ambiguity about the level of BMP expression prior to the MBT; immunofluorescence data suggests that the BMP ligand gradient is formed through downregulation on the ventral side, whereas computational models suggest a combination of ventral upregulation and dorsal downregulation[11], [13], [14]. To account for this ambiguity, we used several modes of ligand gradient formation in the virtual blastula formation simulation. In Figure 5, the top panel illustrates a cartoon representation of ligand expression before and after MBT. The cartoons depict ligand gradient formation by upregulation (Figure 5A), downregulation (Figure 5B), and in a mixed regulation system with ventral upregulation and dorsal downregulation (Figure 5C). In the middle panel, we show the dynamic, minute-wise change in positional information as the receptor tetramer levels respond to changes in the ligand gradient. In all three gradient formation scenarios, the heterodimer-heterotetramer offers competitive positional information transmission under dynamic ligand conditions. In the lower panel of Figure 5, we show the dynamic positional information transmitted via receptor tetramer-induced pSmad signaling. All three scenarios show slight improvement for information transmission via the heterodimer-heterotetramer, allowing it to compete with the other complexes despite occurring at lower levels.

**Figure 5.**
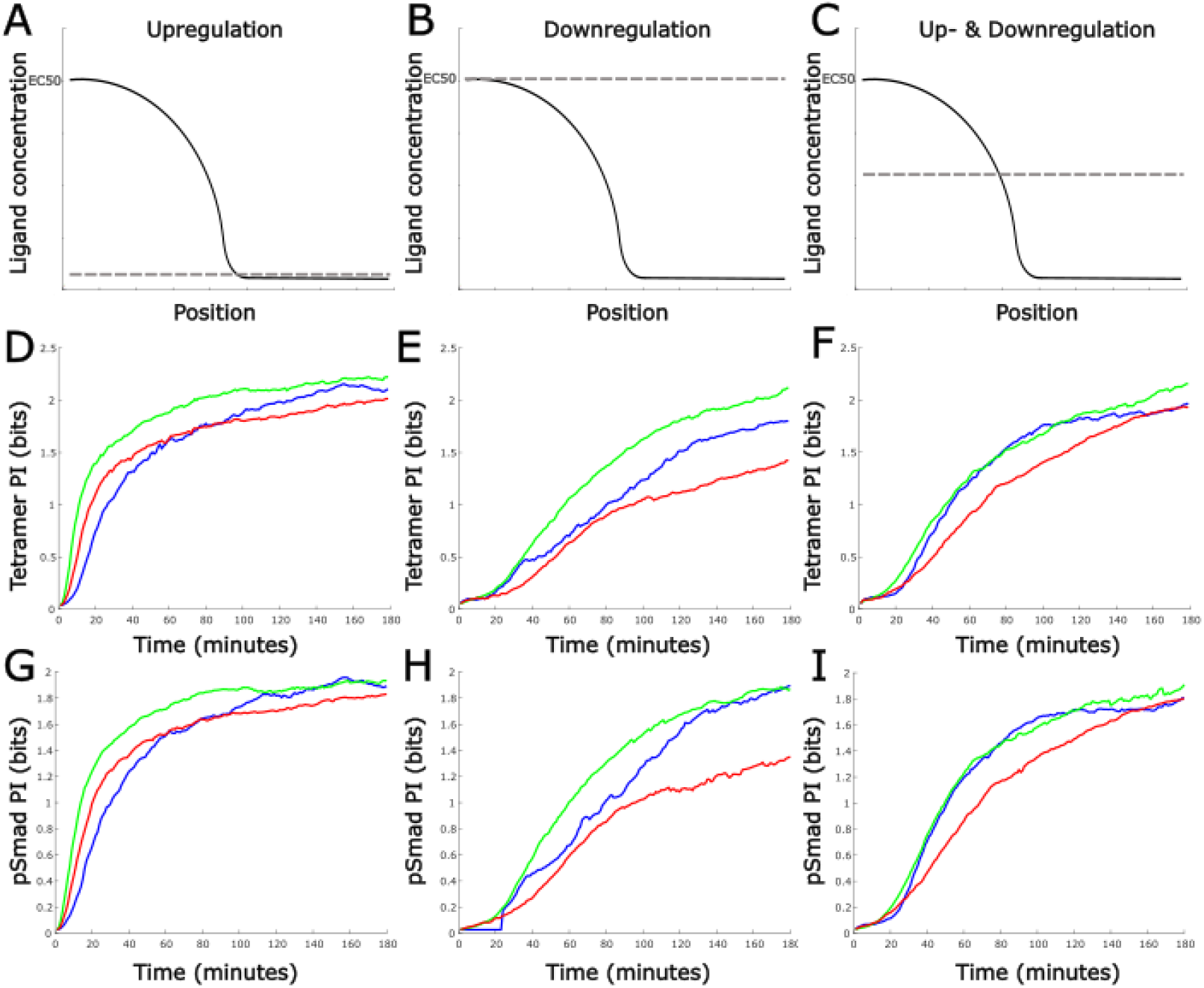
Positional Information transduced by a changing BMP gradient. Positional information is shown from a simulated embryo through three hours of changing gradient simulation, using three regulatory methods. **A-C**. Schematics of the three modes of regulation, each progressing from the dashed line to the final gradient shown by the dark line. **D-F**. Calculated from tetramer levels across the simulated embryo. **G-I**. Calculated from pSmad levels through a simple Smad model acting downstream of receptor formation.

## Discussion & Conclusion

### Heterodimer-heterotetramer strength may arise from a tandem arrangement with pSmad network

The conserved requirement for heterodimer-heteroreceptor signaling in early embryogenesis across evolutionarily distant animal models suggests that this mechanism offers a crucial selective advantage. Our stochastic modeling of BMP ligand-receptor oligomerization reveals that, contrary to our initial hypothesis, heterodimer-heterotetramer oligomerization has greater noise (CV) and less positional information than the other candidate receptor tetramers. However, Fourier analysis reveals that the heterodimer-heterotetramer has relatively less low-frequency noise. Having less low-frequency components manifests as fewer and shorter departures from the mean ± SD. In biological contexts in which downstream autoregulation occurs, long duration departures are particularly maladaptive. Avoiding those long-duration departures during early embryogenesis may be the basis upon which heterodimer-heterotetramer signaling is selected.

Additionally, we find that the integration of pSmad signaling enhances the BMP receptor parameter space for the heterodimer-heterotetramer in terms of positional information transmission. Downstream pSmad signaling may operate as a low-pass filter, which acts to eliminate the high-frequency noise in the heterodimer-heterotetramer signal while the lower-frequency oscillations of homomeric tetramer signals remain unaffected. Interactions with BMP signaling regulators, which are not included in this model, may further filter low-frequency noise, making the information transmission advantages of the heterodimer-heterotetramer more apparent.

### Limitations present in experimental data restrict claims as model results vary greatly over parameter screenings

By simulating the dynamic ligand gradient during MBT, we tested the ability of each receptor tetramer to rapidly interpret changes to the morphogen gradient. We find that the heterodimer-heterotetramer is capable of providing competitive positional information during most of the first three hours of upregulation, downregulation and mixed regulation systems, despite occurring at much lower levels in the system. It remains unknown precisely when fate determination occurs during blastula formation; therefore, either the earlier or later periods of blastula formation may be the critical period. It is also uncertain how rapidly BMP ligand gradient formation occurs after MBT. Further experimental evidence on the temporal profile of BMP ligand gradient formation during and after MBT is needed for critical evaluation of the role of heterodimer-heterotetramer oligomerization in this developmental stage.

The dearth of experimental evidence indicating the absolute ligand levels of the BMP gradient was a significant challenge in constructing our stochastic model. The data from Figure 2A-D suggest that each tetramer receptor has different noise propagation and information transmission properties at different ligand receptor levels. Noise suppression capabilities appear to be linked to the level of receptor saturation; for each receptor tetramer, the coefficient of variation is lowest at ligand levels that are near peak tetramer levels. The heterodimer-heterotetramer requires much higher ligand levels to reach peak tetramer levels and consequently fares better in noise and positional information assessments at higher ligand levels. To overcome our lack of information about the absolute ligand quantity in the embryo, we used a gradient scaled to the EC50 of each tetramer receptor in our stochastic analysis. A similar limitation to our model construction was the lack of information about specific BMP receptor quantities during blastula formation. We also tested the effect of modulating individual BMP receptor levels and found that the heterodimer-heterotetramer experiences an improvement in information transmission in systems with higher ratios of Acvr1 to BmpR1 expression, and this correlation is conversely observed with increasing BMPR1 expression. Experiments designed for absolute quantification of molecular species are useful complements to relative quantification data and may be particularly helpful for further integrating quantitative modeling approaches to developmental biology studies.

### Further analysis required with advanced pSmad filter to confirm noise source in signal transduction pathway

Prior work modeling the BMP system deterministically has suggested that the promiscuity of the interactions between multiple ligand and receptor variants provides for combinatorial signal perception[21], [32]. This complexity makes mechanistic interactions and motivations highly difficult to unravel. There is also a lack of quantified data for major components[27], [33] along with some seemingly-adaptive network reactions, such as the BMP2/2 homodimer having no binding affinity for Acvr1 whatsoever in vitro but being able to rescue embryo signaling when overexpressed in vivo[34]. There may be another non-linearity affecting the system between receptor oligomerization and gene transcription control. More experimental evidence from perturbed systems is necessary to compare with modeling to uncover where unexpected interactions are arising. While many labs are working across multiple systems to track BMP through its downstream signaling network[35], [36], [37], direct measurement of the extracellular morphogen via fixed staining or live imaging has not been successful. Therefore, the noise and attenuation effects in place between BMP and pSmad cannot be uncovered directly through experimentation currently. This gap in system and multiscale modeling of theorized principles can aid in interrogating unexpected results, as well as identifying new experimental directions where unexpected modeling results arise.

Our work is the first to fully model receptor oligomerization in the BMP system. Additionally, our use of stochastic modeling allows us to eschew traditional analysis of non-sequential steady-state data which often fails to capture ‘temporal effects’ and may obscure biologically meaningful results. Our results identify emergent qualities of heterodimer-heterotetramer signaling that arise from the combination of high-affinity and low-affinity interactions between ligand and receptor. Specifically, we identify advantages for the heterodimer-heterotetramer in minimizing error duration and enhancing positional information transfer in dynamic ligand environments. Both properties are particularly advantageous in developmental contexts with tight temporal constraints. Kinetic-based stochastic modeling approaches may offer insight into the role of heterodimer-heterotetramer signaling and, more generally, into developing a systems-level understanding of the role of heterodimer signaling in developmental systems.

## Acknowledgement

This work is based upon efforts supported by EMBRIO Institute, contract #2120200, a National Science Foundation (NSF) Biology Integration Institute.

## Competing Interests

All authors declare no financial or non-financial competing interests.

## Code availability

The underlying code for this study can be accessed via this link https://github.com/larso124/BMP_receptorOligomerization.

## Notes

https://github.com/larso124/BMP_receptorOligomerization

### Competing Interest Statement

The authors have declared no competing interest.

## References

[1] C. Agnew et al., “Structural basis for ALK2/BMPR2 receptor complex signaling through kinase domain oligomerization,” Nat. Commun., vol. 12, no. 1, p. 4950, Aug. 2021, doi: 10.1038/s41467-021-25248-5.

[2] A. M. Turing, “The chemical basis of morphogenesis,” Philos. Trans. R. Soc. Lond. B. Biol. Sci., vol. 237, no. 641, pp. 37–72, Aug. 1952, doi: 10.1098/rstb.1952.0012.

[3] L. Wolpert, “Positional information and the spatial pattern of cellular differentiation,” J. Theor. Biol., vol. 25, no. 1, pp. 1–47, Oct. 1969, doi: 10.1016/S0022-5193(69)80016-0.

[4] J. B. Gurdon and P.-Y. Bourillot, “Morphogen gradient interpretation,” Nature, vol. 413, no. 6858, pp. 797–803, Oct. 2001, doi: 10.1038/35101500.

[5] C. J. Neumann and S. M. Cohen, “Long-range action of Wingless organizes the dorsalventral axis of the Drosophila wing,” Development, vol. 124, no. 4, pp. 871–880, Feb. 1997, doi: 10.1242/dev.124.4.871.

[6] J. A. Tucker, K. A. Mintzer, and M. C. Mullins, “The BMP Signaling Gradient Patterns Dorsoventral Tissues in a Temporally Progressive Manner along the Anteroposterior Axis,” Dev. Cell, vol. 14, no. 1, pp. 108–119, Jan. 2008, doi: 10.1016/j.devcel.2007.11.004.

[7] S. C. Little and M. C. Mullins, “Bone morphogenetic protein heterodimers assemble heteromeric type I receptor complexes to pattern the dorsoventral axis,” Nat. Cell Biol., vol. 11, no. 5, pp. 637–643, May 2009, doi: 10.1038/ncb1870.

[8] S. C. Little and M. C. Mullins, “Extracellular modulation of BMP activity in patterning the dorsoventral axis,” Birth Defects Res. Part C Embryo Today Rev., vol. 78, no. 3, pp. 224–242, Sep. 2006, doi: 10.1002/bdrc.20079.

[9] J. L. Ross, M. Y. Ali, and D. M. Warshaw, “Cargo transport: molecular motors navigate a complex cytoskeleton,” Curr. Opin. Cell Biol., vol. 20, no. 1, pp. 41–47, Feb. 2008, doi: 10.1016/j.ceb.2007.11.006.

[10] J. A. Dutko, “Tyl retromobility is restricted by human APOBEC proteins and facilitated by messenger RNA turnover proteins,” University at Albany, State University of New York, 2008.

[11] J. Zinski, Y. Bu, X. Wang, W. Dou, D. Umulis, and M. C. Mullins, “Systems biology derived source-sink mechanism of BMP gradient formation,” eLife, vol. 6, p. e22199, Aug. 2017, doi: 10.7554/eLife.22199.

[12] S. Tozer, G. L. Dréau, E. Marti, and J. Briscoe, “Temporal control of BMP signalling determines neuronal subtype identity in the dorsal neural tube,” Development, vol. 140, no. 7, pp. 1467–1474, Apr. 2013, doi: 10.1242/dev.090118.

[13] M.-C. Ramel and C. S. Hill, “The ventral to dorsal BMP activity gradient in the early zebrafish embryo is determined by graded expression of BMP ligands,” Dev. Biol., vol. 378, no. 2, pp. 170–182, Jun. 2013, doi: 10.1016/j.ydbio.2013.03.003.

[14] A. P. Pomreinke, G. H. Soh, K. W. Rogers, J. K. Bergmann, A. J. Bläßle, and P. Müller, “Dynamics of BMP signaling and distribution during zebrafish dorsal-ventral patterning,” eLife, vol. 6, p. e25861, Aug. 2017, doi: 10.7554/eLife.25861.

[15] T. Leung et al., “bozozok directly represses bmp2b transcription and mediates the earliest dorsoventral asymmetry of bmp2b expression in zebrafish,” Development, vol. 130, no. 16, pp. 3639–3649, Aug. 2003, doi: 10.1242/dev.00558.

[16] D. S. Koos and R. K. Ho, “The nieuwkoid/dharma Homeobox Gene Is Essential for bmp2b Repression in the Zebrafish Pregastrula,” Dev. Biol., vol. 215, no. 2, pp. 190–207, Nov. 1999, doi: 10.1006/dbio.1999.9479.

[17] T. Shimizu et al., “Cooperative roles of Bozozok/Dharma and Nodal-related proteins in the formation of the dorsal organizer in zebrafish,” Mech. Dev., vol. 91, no. 1–2, pp. 293–303, Mar. 2000, doi: 10.1016/S0925-4773(99)00319-6.

[18] L. Solnica-Krezel and W. Driever, “The role of the homeodomain protein Bozozok in zebrafish axis formation,” Int. J. Dev. Biol., vol. 45, no. 1, pp. 299–310, 2001.

[19] L. Li, X. Wang, M. C. Mullins, and D. M. Umulis, “Evaluation of BMP-mediated patterning in a 3D mathematical model of the zebrafish blastula embryo,” J. Math. Biol., vol. 80, no. 1–2, pp. 505–520, Jan. 2020, doi: 10.1007/s00285-019-01449-x.

[20] L. Li, X. Wang, J. Chai, X. Wang, A. Buganza-Tepole, and D. M. Umulis, “Determining the role of advection in patterning by bone morphogenetic proteins through neural network model-based acceleration of a 3D finite element model of the zebrafish embryo,” Front. Syst. Biol., vol. 2, p. 983372, Oct. 2022, doi: 10.3389/fsysb.2022.983372.

[21] Y. E. Antebi et al., “Combinatorial Signal Perception in the BMP Pathway,” Cell, vol. 170, no. 6, pp. 1184-1196.e24, Sep. 2017, doi: 10.1016/j.cell.2017.08.015.

[22] Y. E. Antebi et al., “Combinatorial Signal Perception in the BMP Pathway,” Cell, vol. 170, no. 6, pp. 1184–1196, Sep. 2017.

[23] M. S. Karim, G. T. Buzzard, and D. M. Umulis, “Secreted, receptor-associated bone morphogenetic protein regulators reduce stochastic noise intrinsic to many extracellular morphogen distributions,” J. R. Soc. Interface, vol. 9, no. 70, pp. 1073–1083, May 2012, doi: 10.1098/rsif.2011.0547.

[24] Md. S. Karim, A. Madamanchi, J. A. Dutko, M. C. Mullins, and D. M. Umulis, “Heterodimer-heterotetramer formation mediates enhanced sensor activity in a biophysical model for BMP signaling,” PLOS Comput. Biol., vol. 17, no. 9, p. e1009422, Sep. 2021, doi: 10.1371/journal.pcbi.1009422.

[25] J. O. Dubuis, G. Tkačik, E. F. Wieschaus, T. Gregor, and W. Bialek, “Positional information, in bits,” Proc. Natl. Acad. Sci., vol. 110, no. 41, pp. 16301–16308, Oct. 2013, doi: 10.1073/pnas.1315642110.

[26] M. J. Thompson, C. A. Young, V. Munnamalai, and D. M. Umulis, “Early radial positional information in the cochlea is optimized by a precise linear BMP gradient and enhanced by SOX2,” Sci. Rep., vol. 13, no. 1, p. 8567, May 2023, doi: 10.1038/s41598-023-34725-4.

[27] L. M. Wakefield, D. M. Smith, T. Masui, C. C. Harris, and M. B. Sporn, “Distribution and modulation of the cellular receptor for transforming growth factor-beta.,” J. Cell Biol., vol. 105, no. 2, pp. 965–975, Aug. 1987, doi: 10.1083/jcb.105.2.965.

[28] T. Menon, A. S. Borbora, R. Kumar, and S. Nair, “Dynamic optima in cell sizes during early development enable normal gastrulation in zebrafish embryos,” Dev. Biol., vol. 468, no. 1–2, pp. 26–40, Dec. 2020, doi: 10.1016/j.ydbio.2020.09.002.

[29] J. H. Abel, B. Drawert, A. Hellander, and L. R. Petzold, “GillesPy: A Python Package for Stochastic Model Building and Simulation,” IEEE Life Sci. Lett., vol. 2, no. 3, pp. 35–38, Sep. 2016, doi: 10.1109/LLS.2017.2652448.

[30] J. E. Ladbury and S. T. Arold, “Noise in cellular signaling pathways: causes and effects,” Trends Biochem. Sci., vol. 37, no. 5, pp. 173–178, May 2012, doi: 10.1016/j.tibs.2012.01.001.

[31] S. Catozzi, J. P. Di-Bella, A. C. Ventura, and J.-A. Sepulchre, “Signaling cascades transmit information downstream and upstream but unlikely simultaneously,” BMC Syst. Biol., vol. 10, no. 1, p. 84, Dec. 2016, doi: 10.1186/s12918-016-0303-2.

[32] K. W. Rogers, M. ElGamacy, B. M. Jordan, and P. Müller, “Optogenetic investigation of BMP target gene expression diversity,” eLife, vol. 9, p. e58641, Nov. 2020, doi: 10.7554/eLife.58641.

[33] J. Zinski, F. Tuazon, Y. Huang, M. Mullins, and D. Umulis, “Imaging and Quantification of P-Smad1/5 in Zebrafish Blastula and Gastrula Embryos,” in Bone Morphogenetic Proteins, vol. 1891, M. B. Rogers, Ed., in Methods in Molecular Biology, vol. 1891., New York, NY: Springer New York, 2019, pp. 135–154. doi: 10.1007/978-1-4939-8904-1_10.

[34] B. Tajer, J. A. Dutko, S. C. Little, and M. C. Mullins, “BMP heterodimers signal via distinct type I receptor class functions,” Proc. Natl. Acad. Sci., vol. 118, no. 15, p. e2017952118, Apr. 2021, doi: 10.1073/pnas.2017952118.

[35] R. M. Monteiro, S. M. C. De Sousa Lopes, M. Bialecka, S. De Boer, A. Zwijsen, and C. L. Mummery, “Real time monitoring of BMP Smads transcriptional activity during mouse development,” genesis, vol. 46, no. 7, pp. 335–346, Jul. 2008, doi: 10.1002/dvg.20402.

[36] R. F. Collery and B. A. Link, “Dynamic smad-mediated BMP signaling revealed through transgenic zebrafish,” Dev. Dyn., vol. 240, no. 3, pp. 712–722, Mar. 2011, doi: 10.1002/dvdy.22567.

[37] H. Al Asafen, A. Beseli, H.-Y. Chen, S. Hiremath, C. M. Williams, and G. T. Reeves, “Dynamics of BMP signaling and stable gene expression in the early Drosophila embryo,” Biol. Open, vol. 13, no. 9, p. bio061646, Sep. 2024, doi: 10.1242/bio.061646.

