## Supplemental for "Stochastic Modeling of BMP Heterodimer-Receptor Interactions Shows Emergence of Low-Pass Filtering Behavior"

**Supplementary Tables**

**Table S1. Table of rate constants**

**Table S2. Model Constraints**

**Table S3. Model Constants, Reactions, and Labels**

**Supplementary Information**

**SI1. Stochastic Simulation**

**SI2. Positional Information**

**Table S1. Table of rate constants**

| Receptor | Units | BMP2/2 | | | BMP7/7 | | | BMP2/7 | | | Ref |
|---|---|---|---|---|---|---|---|---|---|---|---|
| | | Value | Name | $K_D$ (nM) | Value | Name | $K_D$ (nM) | Value | Name | $K_D$ (nM) | |
| $1^{st}$ Alk3/BMPR1 | $k_{on}$: $nM^{-1}s^{-1}$<br>$k_{off}$: $s^{-1}$ | $5 \times 10^{-4}$<br>$4 \times 10^{-4}$ | Ar3on<br>Br3on | 0.8 | $5 \times 10^{-4}$<br>$4 \times 10^{-4}$ | A7r3on<br>A7r3off | 56.4 | *$5 \times 10^{-4}$*<br>*$4 \times 10^{-4}$* | A27r3on<br>A27r3off | 0.8 | 1,2 |
| $2^{nd}$ Alk3/BMPR1 | $k_{on}$: $nM^{-1}s^{-1}$ | $5 \times 10^{-4}$<br>$4 \times 10^{-4}$ | Br3on<br>Br3off | 0.8 | $5 \times 10^{-4}$<br>$4 \times 10^{-4}$ | B7r3on<br>B7r3off | 56.4 | *$1.4 \times 10^{-4}$*<br>*$7.9 \times 10^{-3}$* | B27r3on<br>B27r3off | 56.4 | 1 |

| | | | | | | | | | | | |
|---|---|---|---|---|---|---|---|---|---|---|---|
| | $k_{off}$: $s^{-1}$ | | | | | | | | | | |
| 1st Ak8/Acvr1 | $k_{on}$: $nM^{-1}s^{-1}$ $k_{off}$: $s^{-1}$ | $1.2x10^{-6}$<br>$1.2x10^{-3}$ | Ar8on<br>Ar8off | 1024 | $5x10^{-4}$<br>$4x10^{-4}$ | A7r8on<br>A7r8off | 512 | *$2.3x10^{-6}$*<br>*$1.2x10^{-3}$* | A27r8on<br>A27r8off | 512 | 1 |
| 2nd Alk8/Acvr1 | $k_{on}$: $nM^{-1}s^{-1}$ $k_{off}$: $s^{-1}$ | $1.2x10^{-6}$<br>$1.2x10^{-3}$ | Br8on<br>Br8off | 1024 | $5x10^{-4}$<br>$4x10^{-4}$ | B7r8on<br>B7r8off | 512 | *$1.2x10^{-6}$*<br>*$1.2x10^{-3}$* | B27r8on<br>B27r8off | 1024 | 1 |
| 1st Type II | $k_{on}$: $nM^{-1}s^{-1}$ $k_{off}$: $s^{-1}$ | $1.5x10^{-3}$<br>$7x10^{-2}$ | ArIIon<br>ArIIoff | 46.6 | $5x10^{-4}$<br>$4x10^{-4}$ | A7rIIon<br>A7rIIoff | 6.42 | *$1.4x10^{-3}$*<br>*$9x10^{-3}$* | A27rIIon<br>A27rIIoff | 6.42 | 3 |
| 2nd Type II | $k_{on}$: $nM^{-1}s^{-1}$ $k_{off}$: $s^{-1}$ | $1.5x10^{-3}$<br>$7x10^{-2}$ | BrIIon<br>BrIIoff | 46.6 | $5x10^{-4}$<br>$4x10^{-4}$ | B7rIIon<br>B7rIIoff | 6.42 | *$1.5x10^{-3}$*<br>*$7x10^{-2}$* | B27rIIon<br>B27rIIoff | 46.6 | 3 |

Italicized columns indicate that the values for BMP2/7 interaction with BMP receptors are not directly measured but rather inferred based on homology with BMP2 and BMP7 receptor binding domains.

This table is adapted from “Heterodimer-heterotetramer formation mediates enhanced sensor activity in a biophysical model for BMP signaling,” by Md Shahriar Karim, et al., 2021, *PLOS Computational Biology*, https://doi.org/10.1371/journal.pcbi.1009422.

1. Heinecke, K. et al. Receptor oligomerization and beyond: a case study in bone morphogenetic proteins. BMC Biol 7, 59 (2009).
1. Saremba, S. et al. Type I receptor binding of bone morphogenetic protein 6 is dependent on N-glycosylation of the ligand. FEBS J 275, 172–183 (2008).
2. Kirsch, T., Nickel, J. & Sebald, W. BMP-2 antagonists emerge from alterations in the low affinity binding epitope for receptor BMPR-II. EMBO J 19, 3314–3324 (2000).

**Table S2. Model Constants**

| Constant | Name in code | Value | Description |
|---|---|---|---|
| Boost up | BU | 50 | Enhancement factor that affects reactions constrained to the cell surface |

| | | | |
|---|---|---|---|
| Cell Volume | VOLUME | 2e-13 μm$^3$ | Cell surface area of the zebrafish embryo at 4 hours post fertilization (hpf) is approximately 400 μm$^2$ (Menon et al., 2020). Allowing a thickness of 0.5 μm resulted in the used compartment volume for reactions |
| Stochiometric Constant | STOCH | 8.3e-3 nM/molecule | Conversion factor based on volume and Avogadro Constant |

**Table S3. Model Constants, Reactions, and Labels**

| **Ligand-Receptor Reactions** | | | | | |
|---|---|---|---|---|---|
| | | **BMP2/2 Reactions** | | | |
| **Rxn. No.** | **Reactions** | **GillesPy Rate** | **Value** | **GillesPy Reverse** | **Value Label** |
| 1/2 | BMP2 * Alk3 -> BMP2_Alk3 | k1A | Ar3on * A1 | k1r | Ar3off |
| 3/4 | BMP2 * Alk8 -> BMP2_Alk8 | k2A | Ar8on * A1 | k2r | Ar8off |
| 5/6 | BMP2 * RII -> BMP2_RII | k3A | ArIIon * A1 | k3r | ArIIoff |
| 7/8 | BMP2_Alk3 * Alk3 -> BMP2_Alk3_Alk3 | k4 | Br3on * STOCH * BU | k4r | Br3off |
| 9/10 | BMP2_Alk3 * Alk8 -> BMP2_Alk3_Alk8 | k5 | Ar8on * STOCH * BU | k5r | Ar8off |
| 11/12 | BMP2_Alk3 * RII -> BMP2_Alk3_RII | k6 | ArIIon * STOCH * BU | k6r | ArIIoff |
| 13/14 | BMP2_Alk8 * Alk3 -> BMP2_Alk3_Alk8 | k7 | Ar3on * STOCH * BU | k7r | Ar3off |
| 15/16 | BMP2_Alk8 * Alk8 -> BMP2_Alk8_Alk8 | k8 | Br8on * STOCH * BU | k8r | Br8off |
| 17/18 | BMP2_Alk8 * RII -> BMP2_Alk8_RII | k9 | ArIIon * STOCH * BU | k9r | ArIIoff |
| 19/20 | BMP2_RII * Alk3 -> BMP2_Alk3_RII | k10 | Ar3on * STOCH * BU | k10r | Ar3off |
| 21/22 | BMP2_RII * Alk8 -> BMP2_Alk8_RII | k11 | Ar8on * STOCH * BU | k11r | Ar8off |
| 23/24 | BMP2_RII * RII -> BMP2_RII_RII | k12 | BrIIon * STOCH * BU | k12r | BrIIoff |
| 25/26 | BMP2_Alk8_Alk8 * RII -> BMP2_Alk8_Alk8_RII | k13 | ArIIon * STOCH * BU | k13r | ArIIoff |
| 27/28 | BMP2_RII_RII * Alk3 -> BMP2_Alk3_RII_RII | k14 | Ar3on * STOCH * BU | k14r | Ar3off |
| 29/30 | BMP2_RII_RII * Alk8 -> BMP2_Alk8_RII_RII | k15 | Ar8on * STOCH * BU | k15r | Ar8off |
| 31/32 | BMP2_Alk3_Alk3 * RII -> BMP2_Alk3_Alk3_RII | k16 | ArIIon * STOCH * BU | k16r | ArIIoff |
| 33/34 | BMP2_Alk3_Alk8 * RII -> BMP2_Alk3_Alk8_RII | k17 | ArIIon * STOCH * BU | k17r | ArIIoff |
| 35/36 | BMP2_Alk3_RII * Alk3 -> BMP2_Alk3_Alk3_RII | k18 | Br3on * STOCH * BU | k18r | Br3off |

| | | | | | |
|---|---|---|---|---|---|
| 37/38 | BMP2_Alk3_RII * Alk8 -> BMP2_Alk3_Alk8_RII | k19 | Ar8on * STOCH * BU | k19r | Ar8off |
| 39/40 | BMP2_Alk3_RII * RII -> BMP2_Alk3_RII_RII | k20 | BrIIon * STOCH * BU | k20r | BrIIoff |
| 41/42 | BMP2_Alk8_RII * Alk3 -> BMP2_Alk3_Alk8_RII | k21 | Ar3on * STOCH * BU | k21r | Ar3off |
| 43/44 | BMP2_Alk8_RII * Alk8 -> BMP2_Alk8_Alk8_RII | k22 | Br8on * STOCH * BU | k22r | Br8off |
| 45/46 | BMP2_Alk8_RII * RII -> BMP2_Alk8_RII_RII | k23 | BrIIon * STOCH * BU | k23r | BrIIoff |
| 47/48 | BMP2_Alk3_Alk3_RII * RII -> BMP2_Alk3_Alk3_RII_RII | k24 | BrIIon * STOCH * BU | k24r | BrIIoff |
| 49/50 | BMP2_Alk3_Alk8_RII * RII -> BMP2_Alk3_Alk8_RII_RII | k25 | BrIIon * STOCH * BU | k25r | BrIIoff |
| 51/52 | BMP2_Alk3_RII_RII * Alk3 -> BMP2_Alk3_Alk3_RII_RII | k26 | Br3on * STOCH * BU | k26r | Br3off |
| 53/54 | BMP2_Alk3_RII_RII * Alk8 -> BMP2_Alk3_Alk8_RII_RII | k27 | Ar8on * STOCH * BU | k27r | Ar8off |
| 55/56 | BMP2_Alk8_Alk8_RII * RII -> BMP2_Alk8_Alk8_RII_RII | k28 | BrIIon * STOCH * BU | k28r | BrIIoff |
| 57/58 | BMP2_Alk8_RII_RII * Alk3 -> BMP2_Alk3_Alk8_RII_RII | k29 | Ar3on * STOCH * BU | k29r | Ar3off |
| 59/60 | BMP2_Alk8_RII_RII * Alk8 -> BMP2_Alk8_Alk8_RII_RII | k30 | Br8on * STOCH * BU | k30r | Br8off |
| | | **BMP7/7 Reactions** | | | |
| 61/62 | BMP7 * Alk3 -> BMP7_Alk3 | k31A | A7r3on * A1 | k31r | A7r3off |
| 63/64 | BMP7 * Alk8 -> BMP7_Alk8 | k32A | A7r8on * A1 | k32r | A7r8off |
| 65/66 | BMP7 * RII -> BMP7_RII | k33A | A7rIIon * A1 | k33r | A7rIIoff |
| 67/68 | BMP7_Alk3 * Alk3 -> BMP7_Alk3_Alk3 | k34 | B7r3on * STOCH * BU | k34r | B7r3off |
| 69/70 | BMP7_Alk3 * Alk8 -> BMP7_Alk3_Alk8 | k35 | A7r8on * STOCH * BU | k35r | A7r8off |
| 71/72 | BMP7_Alk3 * RII -> BMP7_Alk3_RII | k36 | A7rIIon * STOCH * BU | k36r | A7rIIoff |
| 73/74 | BMP7_Alk8 * Alk3 -> BMP7_Alk3_Alk8 | k37 | A7r3on * STOCH * BU | k37r | A7r3off |
| 75/76 | BMP7_Alk8 * Alk8 -> BMP7_Alk8_Alk8 | k38 | B7r8on * STOCH * BU | k38r | B7r8off |
| 77/78 | BMP7_Alk8 * RII -> BMP7_Alk8_RII | k39 | A7rIIon * STOCH * BU | k39r | A7rIIoff |
| 79/80 | BMP7_RII * Alk3 -> BMP7_Alk3_RII | k40 | A7r3on * STOCH * BU | k40r | A7r3off |
| 81/82 | BMP7_RII * Alk8 -> BMP7_Alk8_RII | k41 | A7r8on * STOCH * BU | k41r | A7r8off |
| 83/84 | BMP7_RII * RII -> BMP7_RII_RII | k42 | B7rIIon * STOCH * BU | k42r | B7rIIoff |
| 85/86 | BMP7_Alk8_Alk8 * RII -> BMP7_Alk8_Alk8_RII | k43 | A7rIIon * STOCH * BU | k43r | A7rIIoff |
| 87/88 | BMP7_RII_RII * Alk3 -> BMP7_Alk3_RII_RII | k44 | A7r3on * STOCH * BU | k44r | A7r3off |
| 89/90 | BMP7_RII_RII * Alk8 -> BMP7_Alk8_RII_RII | k45 | A7r8on * STOCH * BU | k45r | A7r8off |
| 91/92 | BMP7_Alk3_Alk3 * RII -> BMP7_Alk3_Alk3_RII | k46 | A7rIIon * STOCH * BU | k46r | A7rIIoff |
| 93/94 | BMP7_Alk3_Alk8 * RII -> BMP7_Alk3_Alk8_RII | k47 | A7rIIon * STOCH * BU | k47r | A7rIIoff |
| 95/96 | BMP7_Alk3_RII * Alk3 -> BMP7_Alk3_Alk3_RII | k48 | B7r3on * STOCH * BU | k48r | B7r3off |
| 97/98 | BMP7_Alk3_RII * Alk8 -> BMP7_Alk3_Alk8_RII | k49 | A7r8on * STOCH * BU | k49r | A7r8off |

| | | | | | |
|---|---|---|---|---|---|
| 99/100 | BMP7_Alk3_RII * RII -> BMP7_Alk3_RII_RII | k50 | B7rIIon * STOCH * BU | k50r | B7rIIoff |
| 101/102 | BMP7_Alk8_RII * Alk3 -> BMP7_Alk3_Alk8_RII | k51 | A7r3on * STOCH * BU | k51r | A7r3off |
| 103/104 | BMP7_Alk8_RII * Alk8 -> BMP7_Alk8_Alk8_RII | k52 | B7r8on * STOCH * BU | k52r | B7r8off |
| 105/106 | BMP7_Alk8_RII * RII -> BMP7_Alk8_RII_RII | k53 | B7rIIon * STOCH * BU | k53r | B7rIIoff |
| 107/108 | BMP7_Alk3_Alk3_RII * RII -> BMP7_Alk3_Alk3_RII_RII | k54 | B7rIIon * STOCH * BU | k54r | B7rIIoff |
| 109/110 | BMP7_Alk3_Alk8_RII * RII -> BMP7_Alk3_Alk8_RII_RII | k55 | B7rIIon * STOCH * BU | k55r | B7rIIoff |
| 111/112 | BMP7_Alk3_RII_RII * Alk3 -> BMP7_Alk3_Alk3_RII_RII | k56 | B7r3on * STOCH * BU | k56r | B7r3off |
| 113/114 | BMP7_Alk3_RII_RII * Alk8 -> BMP7_Alk3_Alk8_RII_RII | k57 | A7r8on * STOCH * BU | k57r | A7r8off |
| 115/116 | BMP7_Alk8_Alk8_RII * RII -> BMP7_Alk8_Alk8_RII_RII | k58 | B7rIIon * STOCH * BU | k58r | B7rIIoff |
| 117/118 | BMP7_Alk8_RII_RII * Alk3 -> BMP7_Alk3_Alk8_RII_RII | k59 | A7r3on * STOCH * BU | k59r | A7r3off |
| 119/120 | BMP7_Alk8_RII_RII * Alk8 -> BMP7_Alk8_Alk8_RII_RII | k60 | B7r8on * STOCH * BU | k60r | B7r8off |
| **BMP2/7 Reactions** | | | | | |
| 121/122 | BMP27 * Alk3 -> BMP7_Alk3 | k61A | A27r3on * A1 | k61r | A27r3off |
| 123/124 | BMP27 * Alk8 -> BMP27_Alk8 | k62A | A27r8on * A1 | k62r | A27r8off |
| 125/126 | BMP27 * RII -> BMP27_RII | k63A | A27rIIon * A1 | k63r | A27rIIoff |
| 127/128 | BMP27_Alk3 * Alk3 -> BMP27_Alk3_Alk3 | k64 | B27r3on * STOCH * BU | k64r | B27r3off |
| 129/130 | BMP27_Alk3 * Alk8 -> BMP27_Alk3_Alk8 | k65 | A27r8on * STOCH * BU | k65r | A27r8off |
| 131/132 | BMP27_Alk3 * RII -> BMP27_Alk3_RII | k66 | A27rIIon * STOCH * BU | k66r | A27rIIoff |
| 133/134 | BMP27_Alk8 * Alk3 -> BMP27_Alk3_Alk8 | k67 | A27r3on * STOCH * BU | k67r | A27r3off |
| 135/136 | BMP27_Alk8 * Alk8 -> BMP27_Alk8_Alk8 | k68 | B27r8on * STOCH * BU | k68r | B27r8off |
| 137/138 | BMP27_Alk8 * RII -> BMP27_Alk8_RII | k69 | A27rIIon * STOCH * BU | k69r | A27rIIoff |
| 138/140 | BMP27_RII * Alk3 -> BMP27_Alk3_RII | k70 | A27r3on * STOCH * BU | k70r | A27r3off |
| 141/142 | BMP27_RII * Alk8 -> BMP27_Alk8_RII | k71 | A27r8on * STOCH * BU | k71r | A27r8off |

| | | | | | |
|---|---|---|---|---|---|
| 143/144 | BMP27_RII * RII -> BMP27_RII_RII | k72 | B27rIIon * STOCH * BU | k72r | B27rIIoff |
| 145/146 | BMP27_Alk8_Alk8 * RII -> BMP27_Alk8_Alk8_RII | k73 | A27rIIon * STOCH * BU | k73r | A27rIIoff |
| 147/148 | BMP27_RII_RII * Alk3 -> BMP27_Alk3_RII_RII | k74 | A27r3on * STOCH * BU | k74r | A27r3off |
| 149/150 | BMP27_RII_RII * Alk8 -> BMP27_Alk8_RII_RII | k75 | A27r8on * STOCH * BU | k75r | A27r8off |
| 151/152 | BMP27_Alk3_Alk3 * RII -> BMP27_Alk3_Alk3_RII | k76 | A27rIIon * STOCH * BU | k76r | A27rIIoff |
| 153/154 | BMP27_Alk3_Alk8 * RII -> BMP27_Alk3_Alk8_RII | k77 | A27rIIon * STOCH * BU | k77r | A27rIIoff |
| 155/156 | BMP27_Alk3_RII * Alk3 -> BMP27_Alk3_Alk3_RII | k78 | B27r3on * STOCH * BU | k78r | B27r3off |
| 157/158 | BMP27_Alk3_RII * Alk8 -> BMP27_Alk3_Alk8_RII | k79 | A27r8on * STOCH * BU | k79r | A27r8off |
| 159/160 | BMP27_Alk3_RII * RII -> BMP27_Alk3_RII_RII | k80 | B27rIIon * STOCH * BU | k80r | B27rIIoff |
| 161/162 | BMP27_Alk8_RII * Alk3 -> BMP27_Alk3_Alk8_RII | k81 | A27r3on * STOCH * BU | k81r | A27r3off |
| 163/164 | BMP27_Alk8_RII * Alk8 -> BMP27_Alk8_Alk8_RII | k82 | B27r8on * STOCH * BU | k82r | B27r8off |
| 165/166 | BMP27_Alk8_RII * RII -> BMP27_Alk8_RII_RII | k83 | B27rIIon * STOCH * BU | k83r | B27rIIoff |
| 167/168 | BMP27_Alk3_Alk3_RII * RII -> BMP27_Alk3_Alk3_RII_RII | k84 | B27rIIon * STOCH * BU | k84r | B27rIIoff |
| 169/170 | BMP27_Alk3_Alk8_RII * RII -> BMP27_Alk3_Alk8_RII_RII | k85 | B27rIIon * STOCH * BU | k85r | B27rIIoff |
| 171/172 | BMP27_Alk3_RII_RII * Alk3 -> BMP27_Alk3_Alk3_RII_RII | k86 | B27r3on * STOCH * BU | k86r | B27r3off |
| 173/174 | BMP27_Alk3_RII_RII * Alk8 -> BMP27_Alk3_Alk8_RII_RII | k87 | A27r8on * STOCH * BU | k87r | A27r8off |
| 175/176 | BMP27_Alk8_Alk8_RII * RII -> BMP27_Alk8_Alk8_RII_RII | k88 | B27rIIon * STOCH * BU | k88r | B27rIIoff |

| | | | | | |
|---|---|---|---|---|---|
| 177/178 | BMP27_Alk8_RII_RII * Alk3 -> BMP27_Alk3_Alk8_RII_RII | k89 | A27r3on * STOCH * BU | k89r | A27r3off |
| 179/180 | BMP27_Alk8_RII_RII * Alk8 -> BMP27_Alk8_Alk8_RII_RII | k90 | B27r8on * STOCH * BU | k90r | B27r8off |
| **Endocytosis Reactions** | | | | | |
| **Rxn. No.** | **Reactions** | **GillesPy Rate** | **Value** | **Name of Reaction** | |
| 1001 | BMP2_Alk3 -> Alk3 | k1000 | 0.0005 | endo1 | |
| 1002 | BMP2_Alk8 -> Alk8 | k1000 | 0.0005 | endo2 | |
| 1003 | BMP2_RII -> RII | k1000 | 0.0005 | endo3 | |
| 1004 | BMP2_Alk3_Alk3 -> Alk3 * Alk3 | k1000 | 0.0005 | endo4 | |
| 1005 | BMP2_Alk3_Alk8 -> Alk3 * Alk8 | k1000 | 0.0005 | endo5 | |
| 1006 | BMP2_Alk3_RII -> Alk3 * RII | k1000 | 0.0005 | endo6 | |
| 1007 | BMP2_Alk8_Alk8 -> Alk8 * Alk8 | k1000 | 0.0005 | endo7 | |
| 1008 | BMP2_Alk8_RII -> Alk8 * RII | k1000 | 0.0005 | endo8 | |
| 1009 | BMP2_RII_RII -> RII * RII | k1000 | 0.0005 | endo9 | |
| 1010 | BMP2_Alk3_Alk3_RII -> Alk3 * Alk3 * RII | k1000 | 0.0005 | endo10 | |
| 1011 | BMP2_Alk3_Alk8_RII -> Alk3 * Alk8 * RII | k1000 | 0.0005 | endo11 | |
| 1012 | BMP2_Alk3_RII_RII -> Alk3 * RII * RII | k1000 | 0.0005 | endo12 | |
| 1013 | BMP2_Alk8_Alk8_RII -> Alk8 * Alk8 * RII | k1000 | 0.0005 | endo13 | |
| 1014 | BMP2_Alk8_RII_RII -> Alk8 * RII * RII | k1000 | 0.0005 | endo14 | |
| 1015 | BMP2_Alk3_Alk3_RII_RII -> Alk3 * Alk3 * RII * RII | k1000 | 0.0005 | endo15 | |
| 1016 | BMP2_Alk3_Alk8_RII_RII -> Alk3 * Alk8 * RII * RII | k1000 | 0.0005 | endo16 | |
| 1017 | BMP2_Alk8_Alk8_RII_RII -> Alk8 * Alk8 * RII * RII | k1000 | 0.0005 | endo17 | |
| 7001 | BMP7_Alk3 -> Alk3 | k7000 | 0.0005 | endo71 | |
| 7002 | BMP7_Alk8 -> Alk8 | k7000 | 0.0005 | endo72 | |
| 7003 | BMP7_RII -> RII | k7000 | 0.0005 | endo73 | |
| 7004 | BMP7_Alk3_Alk3 -> Alk3 * Alk3 | k7000 | 0.0005 | endo74 | |
| 7005 | BMP7_Alk3_Alk8 -> Alk3 * Alk8 | k7000 | 0.0005 | endo75 | |
| 7006 | BMP7_Alk3_RII -> Alk3 * RII | k7000 | 0.0005 | endo76 | |
| 7007 | BMP7_Alk8_Alk8 -> Alk8 * Alk8 | k7000 | 0.0005 | endo77 | |
| 7008 | BMP7_Alk8_RII -> Alk8 * RII | k7000 | 0.0005 | endo78 | |

| | | | | |
|---|---|---|---|---|
| 7009 | BMP7_RII_RII -> RII * RII | k7000 | 0.0005 | endo79 |
| 7010 | BMP7_Alk3_Alk3_RII -> Alk3 * Alk3 * RII | k7000 | 0.0005 | endo710 |
| 7011 | BMP7_Alk3_Alk8_RII -> Alk3 * Alk8 * RII | k7000 | 0.0005 | endo711 |
| 7012 | BMP7_Alk3_RII_RII -> Alk3 * RII * RII | k7000 | 0.0005 | endo712 |
| 7013 | BMP7_Alk8_Alk8_RII -> Alk8 * Alk8 * RII | k7000 | 0.0005 | endo713 |
| 7014 | BMP7_Alk8_RII_RII -> Alk8 * RII * RII | k7000 | 0.0005 | endo714 |
| 7015 | BMP7_Alk3_Alk3_RII_RII -> Alk3 * Alk3 * RII * RII | k7000 | 0.0005 | endo715 |
| 7016 | BMP7_Alk3_Alk8_RII_RII -> Alk3 * Alk8 * RII * RII | k7000 | 0.0005 | endo716 |
| 7017 | BMP7_Alk8_Alk8_RII_RII -> Alk8 * Alk8 * RII * RII | k7000 | 0.0005 | endo717 |
| 2001 | BMP27_Alk3 -> Alk3 | k2000 | 0.0005 | endo271 |
| 2002 | BMP27_Alk8 -> Alk8 | k2000 | 0.0005 | endo272 |
| 2003 | BMP27_RII -> RII | k2000 | 0.0005 | endo273 |
| 2004 | BMP27_Alk3_Alk3 -> Alk3 * Alk3 | k2000 | 0.0005 | endo274 |
| 2005 | BMP27_Alk3_Alk8 -> Alk3 * Alk8 | k2000 | 0.0005 | endo275 |
| 2006 | BMP27_Alk3_RII -> Alk3 * RII | k2000 | 0.0005 | endo276 |
| 2007 | BMP27_Alk8_Alk8 -> Alk8 * Alk8 | k2000 | 0.0005 | endo277 |
| 2008 | BMP27_Alk8_RII -> Alk8 * RII | k2000 | 0.0005 | endo278 |
| 2009 | BMP27_RII_RII -> RII * RII | k2000 | 0.0005 | endo279 |
| 2010 | BMP27_Alk3_Alk3_RII -> Alk3 * Alk3 * RII | k2000 | 0.0005 | endo2710 |
| 2011 | BMP27_Alk3_Alk8_RII -> Alk3 * Alk8 * RII | k2000 | 0.0005 | endo2711 |
| 2012 | BMP27_Alk3_RII_RII -> Alk3 * RII * RII | k2000 | 0.0005 | endo2712 |
| 2013 | BMP27_Alk8_Alk8_RII -> Alk8 * Alk8 * RII | k2000 | 0.0005 | endo2713 |
| 2014 | BMP27_Alk8_RII_RII -> Alk8 * RII * RII | k2000 | 0.0005 | endo2714 |
| 2015 | BMP27_Alk3_Alk3_RII_RII -> Alk3 * Alk3 * RII * RII | k2000 | 0.0005 | endo2715 |
| 2016 | BMP27_Alk3_Alk8_RII_RII -> Alk3 * Alk8 * RII * RII | k2000 | 0.0005 | endo2716 |
| 2017 | BMP27_Alk8_Alk8_RII_RII -> Alk8 * Alk8 * RII * RII | k2000 | 0.0005 | endo2717 |

### SI1. Stochastic Simulation

A GillesPy2 model contains species, parameters, and reactions. All these objects are contained in table S3, with the species being the components in the reactions – except for the BMP ligands, BMP2, BMP7, and BMP27, which are not created as specific species but included in reactions with their effective concentrations adjusting the reaction rates.

The SSACSolver uses the 'direct method' to calculate samples from the chemical master equation (CME), utilizing Monte Carlo simulation (Gillespie et al., 2013; Matthew et al., 2023). The CME is reprinted here for reference.

$$\frac{\partial P(X,t)}{\partial t}\sum_{j=1}^{M}\left(a_j(X-v_j)P\left(X-v_j,t\middle|X_0,t_0\right)-a_j(X)P(X,t|X_0,t_0)\right)$$

The joint probability density function (PDF) is used to generate random timestep τ and random molecular interaction j (part of the Michaelis-Menten reaction set), which determine simulation steps and states. The PDF is reprinted here for reference.

$$\Pr\,(\tau,j|X,t)=a_j(X)e^{-\sum_{i=1}^{M}a_i(X)\tau}$$

This system is memoryless, and determining a next state is not dependent on anything that has occurred in the system prior to the current state. These calculations are encoded within the GillesPy2 package and are implemented on the system of equations in table S3, using values from tables S1 and S2.

### SI2. Positional Information

Positional information calculations were guided by Matt Thompson (Thompson et al., 2023), based on work from Tkacik et al. in quantifying information provided to cells through a spatial profile (Tkačik et al., 2015).

Calculation were completed using the 'direct method', with binning(Δ, [2:6:20]) and bootstrapping(M, [0.05, 0.75, 0.80, 0.85, 0.90, 0.95]). The equation is reprinted here for reference.

$$I_{\Delta,M}(\{\mathrm{g}_i\};\mathrm{x})=\sum_{\{\mathrm{g}_i\};\mathrm{x}}\tilde{P}_{\Delta,M}(\{\mathrm{g}_i\};\mathrm{x})\log_2\frac{\tilde{P}_{\Delta,M}(\{\mathrm{g}_i\};\mathrm{x})}{\tilde{P}_{x\Delta,M}(\{\mathrm{g}_i\};\mathrm{x})\tilde{P}_{\Delta,M}(\{\{\mathrm{g}_i\}\})}$$

DOI: 10.1534/genetics.114.171850

The summation results in a naïve estimate of the information provided by a profile/gradient (g) across the evenly distributed nuclei (with position x). Extrapolation to very small bin sizes gives a direct estimate of the positional information.

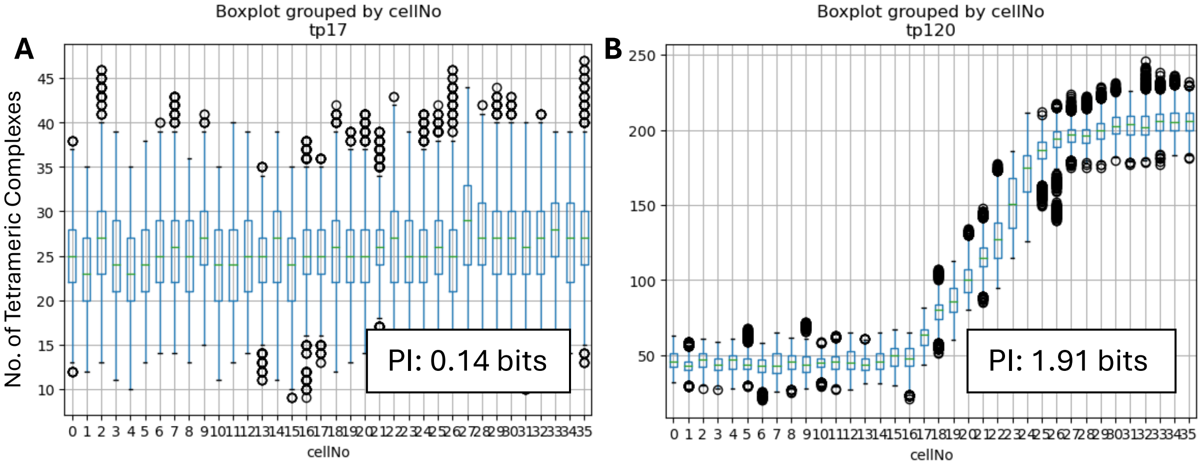

**Figure S1. Progression of positional information provided by tetrameric complexes across an embryo.**

Positional information (PI) was calculated across 30 runs of the simulation, using the output levels of the heterodimer-heterotetramer complex as the gradient across 36 simulated cells. A) Timepoint (minute) 17 of the three-hour simulation gives 0.14 bits of information. B) Once the gradient has had 120 minutes to develop, the positional information has increased to 1.91 bits. These samples are taken from the mid-regulation method of gradient development.
